# Multi-tissue polygenic models for transcriptome-wide association studies

**DOI:** 10.1101/107623

**Authors:** Yongjin Park, Abhishek Sarkar, Kunal Bhutani, Manolis Kellis

## Abstract

Transcriptome-wide association studies (TWAS) have proven to be a powerful tool to identify genes associated with human diseases by aggregating cis-regulatory effects on gene expression. However, TWAS relies on building predictive models of gene expression, which are sensitive to the sample size and tissue on which they are trained. The Gene Tissue Expression Project has produced reference transcriptomes across 53 human tissues and cell types; however, the data is highly sparse, making it difficult to build polygenic models in relevant tissues for TWAS. Here, we propose fQTL, a multi-tissue, multivariate model for mapping expression quantitative trait loci and predicting gene expression. Our model decomposes eQTL effects into SNP-specific and tissue-specific components, pooling information across relevant tissues to effectively boost sample sizes. In simulation, we demonstrate that our multi-tissue approach outperforms single-tissue approaches in identifying causal eQTLs and tissues of action. Using our method, we fit polygenic models for 13,461 genes, characterized the tissue-specificity of the learned *cis*-eQTLs, and performed TWAS for Alzheimer’s disease and schizophrenia, identifying 107 and 382 associated genes, respectively.

## II. INTRODUCTION

A fundamental barrier to interpreting the role of non-coding genetic variation identified by genome-wide association studies (GWAS) is understanding how specific single nucleotide polymorphisms (SNPs) cause changes in gene expression, and how those changes in expression lead to downstream phenotypes. Recently, transcriptome wide association studies (TWAS) have proven to be a powerful tool to predict the impact of cis-regulatory regions on expression and directly associate genes with downstream disease phenotypes^1,2^. The key idea of TWAS is to train multivariate expression quantitative trait loci (eQTL) models on reference expression panels, use these models to predict (impute) unobserved gene expression in large scale GWAS cohorts, and compute association statistics by regressing phenotype directly onto imputed gene expression.

The success of TWAS is dependent on the sample size of the expression reference panel and the tissue in which expression was measured. The Gene Tissue Expression (GTEx) Project has produced reference expression profiles for 450 individuals across 53 tissues; however, only a subset of individuals were measured for each tissue^3^. The GTEx dataset is highly sparse: 70% of individual-tissue-gene expression observations are missing. Here, we seek to improve polygenic eQTL modeling by pooling information across tissues to both overcome limited sample size and improve prediction accuracy.

Previously proposed multi-tissue (multi-trait) QTL approaches have primarily focused on identifying the tissue of action using Bayesian model comparison^4,5^. However, combinatorial searches over models quickly become intractable; for example, to find the best model for 20 tissues, we need to compare millions of candidate models for each gene. Moreover, existing approaches still assume independence of SNPs and train univariate eQTL models. In parallel, several multivariate regression methods have been proposed to identify causal eQTLs within single tissues, based either on forward selection^6^ or Bayesian variable selection^7 –9^.

Here, we propose factored polygenic QTL analysis (fQTL), a multi-response, multivariate regression model for eQTL mapping, to jointly identify causal eQTLs and tissues of action. Our model has several advantages over existing methods: (1) We fit a multivariate regression model jointly modeling the effect of all *cis*-SNPs on expression while performing variable selection over SNPs in linkage disequilibrium (LD). (2) **W**e jointly model multiple gene expression vectors across tissues within a single unified model to pool information across relevant tissues. (3) We factorize the SNP by tissue effect size matrix into genomic location-dependent and tissue-dependent components, improving the interpretability of the fitted model. We demonstrate that our approach outperforms existing methods in a variety of realistic simulations. We trained fQTL models for 13,641 genes on the GTEx reference cohort, used them to characterize multi-tissue regulatory contexts in TWAS analysis of Alzheimer’s disease (AD) and schizophrenia (SCZ), and identified 107 AD-associated and 382 SCZ-associated genes. Several of the genes provide interesting clues to disease mechanisms and should be further investigated in follow-up studies. **W**e have made software implementing the method, trained model weights, and full TWAS summary statistics publicly available (https://github.com/ypark/fqtl).

## III. METHODS

### A. Factored QTL model specification

We propose `fQTL`, a multi-response, multivariate regression approach to jointly model gene expression across tissues and individuals. For each gene, we assume the columns of the *n* × *m* (individual-by-tissue) gene expression matrix *Y* are normalized to be approximately Gaussian. We regress *Y* onto the *n* × *p* genotype matrix *X*, which is centered (but not scaled). We fit a linear model:

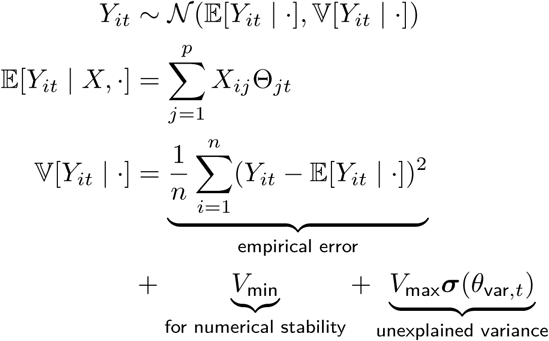

We define Θ_*jt*_ to be the effect of SNP *j* on gene expression of that gene in tissue *t*. We define σ(*z*) = 1/(1 + exp(−*z*)) and we set 
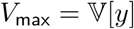
 and *V*_min_ = 10^−4^*V*_max_.

The key idea of fQTL is that we assume the eQTL effect size matrix Θ can be decomposed into tissue-invariant components *θ*^snp^ and tissue-dependent components *θ*^tis^:

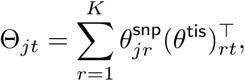

Here, we assume *K* = 1 for ease of interpreting the results, although our inference algorithm (described below) supports fitting arbitrary *K* ≤ *m*.

The fundamental problem in fitting a multivariate regression on genotypes *X* is co-linearity of the columns of *X* due to LD. Co-linearity causes ordinary regression to be ill-posed (infinitely many solutions), but a Bayesian regression with the spike and slab prior can successfully solve the variable selection problem^10    –15^. Here, we assume the spike and slab (point-normal mixture) prior on each column of *θ*^snp^ and *θ*^tis^. Considering one of the columns *θ*:

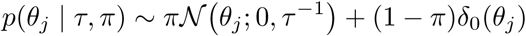

where 𝒩(*x*; ·, ·) denotes the Gaussian density and *δ*(*x*) denotes the Dirac delta function (point mass at zero).

### B. Stochastic variational inference

For our model, the spike and slab prior is non-conjugate to the likelihood. Therefore, estimating the posterior distribution of *θ* requires either Markov Chain Monte Carlo (MCMC)^16,17^, expectation propagation^18,19^, or variational inference^13,20^. Based on our preliminary experiments, we chose not to use MCMC due to convergence problems. Instead, we perform variational inference: we re-cast the problem as an optimization problem over variational parameters ψ = (*α*, *β*, *γ*) finding the best mean-field approximating distribution *q*(*θ* | ψ) to the intractable posterior *p*(*θ* | *X*, *Y*).

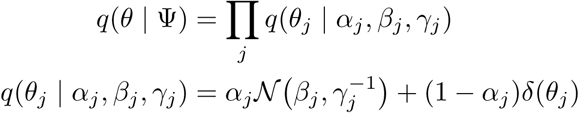

Under the variational approximation, the parameters *θ_j_* are mutually independent and their means and variances (approximate posterior means and posterior variances) can be written in terms of the variational parameters:

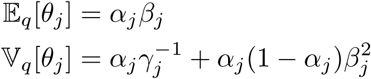

Our goal is to find the *q* with minimum KL-divergence with *p* by optimizing over ψ; however, in general the KL-divergence does not have an analytic form. Instead, we solve an equivalent optimization problem, maximizing the evidence lower-bound (ELBO):

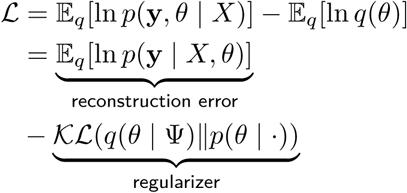

Prior work derived coordinate ascent updates to maximize the ELBO assuming hyperparameters (*π*, *τ*, *σ*^2^) were fixed^13^. The method is scalable due to fast convergence, but is sensitive to the hyperparameters and to the initialization of the variational parameters. In practice, the algorithm requires two outer loops of importance sampling steps over the hyperparameter space: one to find an optimal initialization, and one to actually perform the optimization from a warm start.

We instead used stochastic optimization to optimize the variational objective, an approach known as stochastic variational inference (SVI)^21,22^. The key idea is to perform gradient ascent, taking steps along the natural gradient of the objective function^23^ with respect to the variational parameters:

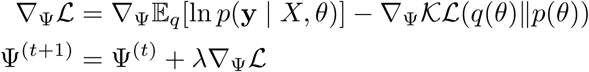

For our choice of approximating family *q*, the regularizer term above has an analytic form, as previously derived^13^:

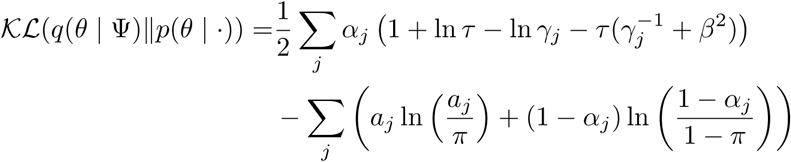

Although we can analytically take gradients of the regularizer (not shown), our use of the spike and slab prior means it is not possible to take gradients of 
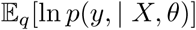
. We use the log-derivative trick to obtain an unbiased estimator for the natural gradients (see Supplementary Methods)^21^:

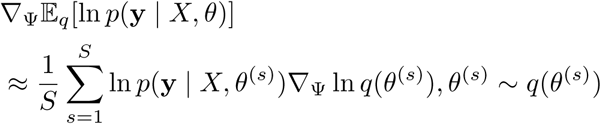

Naive implementation of SVI is slow to converge because of the variance of the stochastic gradient. To speed up convergence, we used a variance reduction technique known as control variates^21,22^, developing a novel control variate for our problem (see Supplementary Methods). We additionally re-parameterized the model in terms of the genetic values *η* = *Xθ*, a parameterization which has been previously studied^24,25.^ This parameterization has several advantages: (1) We avoid sampling p-vectors *θ*, reducing the computation time for *n* < *p* (which is true in our setting). (2) **W**e further reduce the variance of the stochastic gradient^25^.

Even with variance reduction techniques, the convergence rate and sparsity of the final fitted model is highly sensitive to the hyperparameters and learning rate. We developed a variational approximation for the hyperparameters (see Supplementary Methods) which allowed us to simultaneously fit the model parameters *θ* and hyperparameters (π, τ). We tuned the learning rate on simulated datasets, finding that a fixed value 0.01 worked well.

### C. Factored QTL model inference

In order to fit fQTL using SVI, we only need to characterize the distribution of *η* under the variational approximation, relying on a previously derived result for the variance of the product of two random vectors^26^:

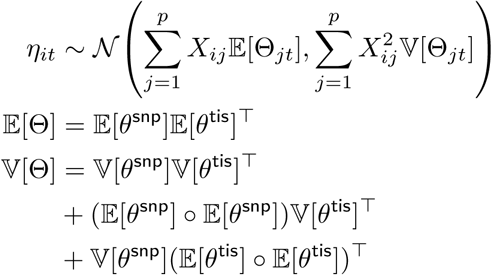

where ◦ denotes elementwise multiplication.

### D. Simulation study

We simulated multi-tissue gene expression matrices *Y* using genotypes in 450 individuals from the specified *cis*-regulatory window around each gene, varying the number of causal eQTLs, the number of tissues of action, and the proportion of variance of gene expression *h^2^* explained by SNP-specific effects. For each setting, we simulated 175 randomly chosen genes. We matched the number of missing observations in each tissue to the number of missing observations in each GTEx tissue. We then quantile-normalized the generated expression values to match real GTEx expression profiles.

We sampled SNP-specific effects 
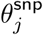
 and tissue-specific effects 
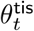
 from the standard normal distribution. We then generated gene expression within tissues with non-zero tissue effect (i.e., tissues within which gene expression could also be genetically controlled) using a linear model^17,27^:

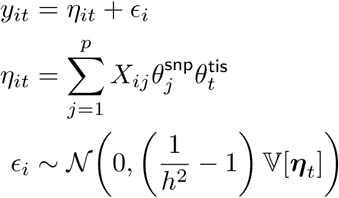

For tissues with zero tissue effect, we sampled expression values from 
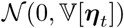
.

### E. Identification of tissues of action in GTEx

We obtained individual genotypes and gene-level RNA-seq read counts from the GTEx consortium (version 6). We restricted our analysis to coding genes annotated by GENCODE v19 and tissues with sample size *n* ≥ 50. Within each tissue, we further restricted to genes having read count greater than 10 in at least 50% of samples. To convert read counts to logarithmic scale, rlog transformed the data using the DESeq2 package^28^. We confirmed after the rlog transformation there was no mean-variance correlation.

For GTEx gene expression, significant PVE is attributable to experimental and technical confounding variables. For each tissue, we jointly fit a polygenic QTL model and performed sparse matrix factorization of the residual to identify non-genetic, technical confounders on the 1000 most variable genes in each tissue. We modeled the *k*-th most variable gene’s expression in *i*-th sample as:

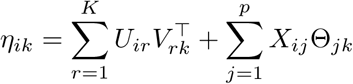

Here, the matrix of genetic effects Θ_*jk*_ denotes the effect of SNP *j* on gene *k*. We fixed Θ_*jk*_ = 0 if SNP *j* was not located within the specified *cis*-regulatory window of gene *k*. We enforced rank sparsity of the factorization by assuming a group spike and slab prior^18^ on *U* and *V* (see Supplementary Methods). We defined learned covariates as columns of *U* with column posterior inclusion probability PIP > 0.5. We included these learned covariates as well as known covariates in subsequent analysis by introducing another linear term in our original model:

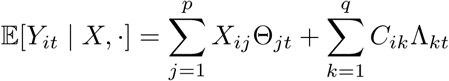

To account for correlations between the columns of *C*, we assume the spike and slab prior on the elements Λ_*kt*_, and incorporate the entire matrix into our stochastic variational inference algorithm described above.

### F. TWAS using factored QTL effects

We perform tissue-specific TWAS by defining tissue-specific QTL effects 
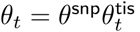
. Prior work proposed a summary-based TWAS statistic 
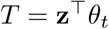
 and characterized its null distribution1. However, the proposed statistic depends only on a point estimate of *θ_t_*, while our factored QTL model estimates both a posterior mean and a posterior variance of *θ_t_*.

We derive the null distribution of *T*, accounting for the posterior variance of the estimated effects. As before, we assume the null distribution of GWAS z-scores follows the multivariate Gaussian distribution, 
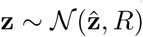
 where the covariance matrix 
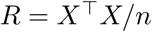
 is estimated from the GWAS cohort genotypes (or approximated using a reference cohort).

Using previously derived results for the mean and variance of the inner product of two stochastic vectors^26^, we can define 
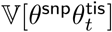
, and use the same result to define 
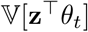
.

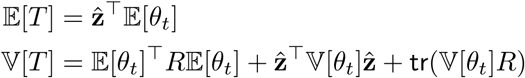

Compared to the previously derived null distribution, the null distribution of our TWAS statistic has two additional terms in the variance which additionally penalize the model complexity of the polygenic QTL model. This modified variance leads to a more conservative hypothesis test.

We performed summary-based tissue-specific TWAS using our method for Alzheimer’s disease and schizophrenia. We downloaded summary statistics for Alzheimer’s disease from the International Genomics of Alzheimer’s Project (http://web.pasteur-lille.fr/en/recherche/u744/igap/igap_download.php) and statistics for schizophrenia from the Psychiatric Genetics Consortium (http://www.med.unc.edu/pgc).

International Genomics of Alzheimer’s Project (IGAP) is a large two-stage study based upon genome-wide association studies (GWAS) on individuals of European ancestry. In stage 1, IGAP used genotyped and imputed data on 7,055,881 single nucleotide polymorphisms (SNPs) to meta-analyze four previously-published GWAS datasets consisting of 17,008 Alzheimer’s disease cases and 37,154 controls (The European Alzheimer’s disease Initiative – EADI the Alzheimer Disease Genetics Consortium – ADGC The Cohorts for Heart and Aging Research in Genomic Epidemiology consortium – CHARGE The Genetic and Environmental Risk in AD consortium – GERAD). In stage 2, 11,632 SNPs were genotyped and tested for association in an independent set of 8,572 Alzheimer’s disease cases and 11,312 controls. Finally, a meta-analysis was performed combining results from stages 1 & 2.

## IV. RESULTS

### A. Factored QTL model specification and inference

We propose fQTL, a multi-response, multivariate regression approach to jointly model gene expression variation across individuals and tissues. For each gene, we fit a linear model regressing expression in individuals *i* and tissues *t* against *cis*-SNPs *j*:

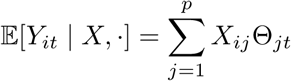

The key idea of fQTL is to assume the eQTL effect size matrix Θ can be decomposed into tissue-invariant components *θ*snp and tissue-dependent components *θ*^tis^:

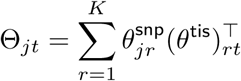

In this study, we focus on *K* = 1 for intuitive interpretation, although our method supports arbitrary *K*. In this case, the intuition behind the factorization is that the estimated effect of a specific SNP on gene expression in a specific tissue could be explained by several possible mechanisms. (1) The variant might disrupt a regulatory motif and alter transcription factor binding, altering gene expression. (2) The region harboring the variant might not be active in the specific tissue, due to reduced chromatin accessibility or altered epigenomic modifications. (3) The upstream transcription factor which mediates the effect of the SNP might be differentially expressed in the specific tissue. Explanation (1) implies the effect is invariant across tissues since it is determined solely by sequence, while explanations (2) and (3) can be tissue-specific. By factorizing the matrix of effects Θ, we allow our model to fit both types of effects. As *K* increases, we deposit the existence of different causal eQTLs for different subsets of tissues, but do not explore this possibility in this study.

The challenge in fitting polygenic QTL models is that the columns of *X* are co-linear due to LD, and we need to select a sparse set of relevant predictors using the spike and slab prior^13^. This prior is non-conjugate to the likelihood, preventing its widespread use in polygenic modeling. We developed a general computational framework to efficiently perform approximate Bayesian inference with this prior in large-scale sparse regression models. The key ideas of our approach are: (1) We use techniques from from black-box variational inference to allow rapid model specification and implementation^22^. (2) We develop novel variance reduction techniques to improve stochastic variational inference for these models^21^. (3) We use our framework to implement sparse regression and matrix factorization, which are the building blocks of fQTL. We have made C++ source code implementing the framework and an R package wrapping the models publicly available (https://github.com/ypark/fqtl).

Our analysis pipeline is shown in Fig. 1. For each tissue, we jointly performed matrix factorization and multi-response (across genes rather than across tissues), multivariate *cis*-eQTL mapping to learn non-genetic covariates on the 1,000 most variable genes. We then corrected both known and learned covariates in all genes and fit our multi-tissue fQTL model including SNPs from 500kb upstream of the transcription start 5 site to 500kb downstream of the transcription end site. Finally, we used the estimated SNP and tissue effects to perform tissue-specific TWAS, accounting for the posterior variance of the estimated QTL effects.

**Figure 1:**
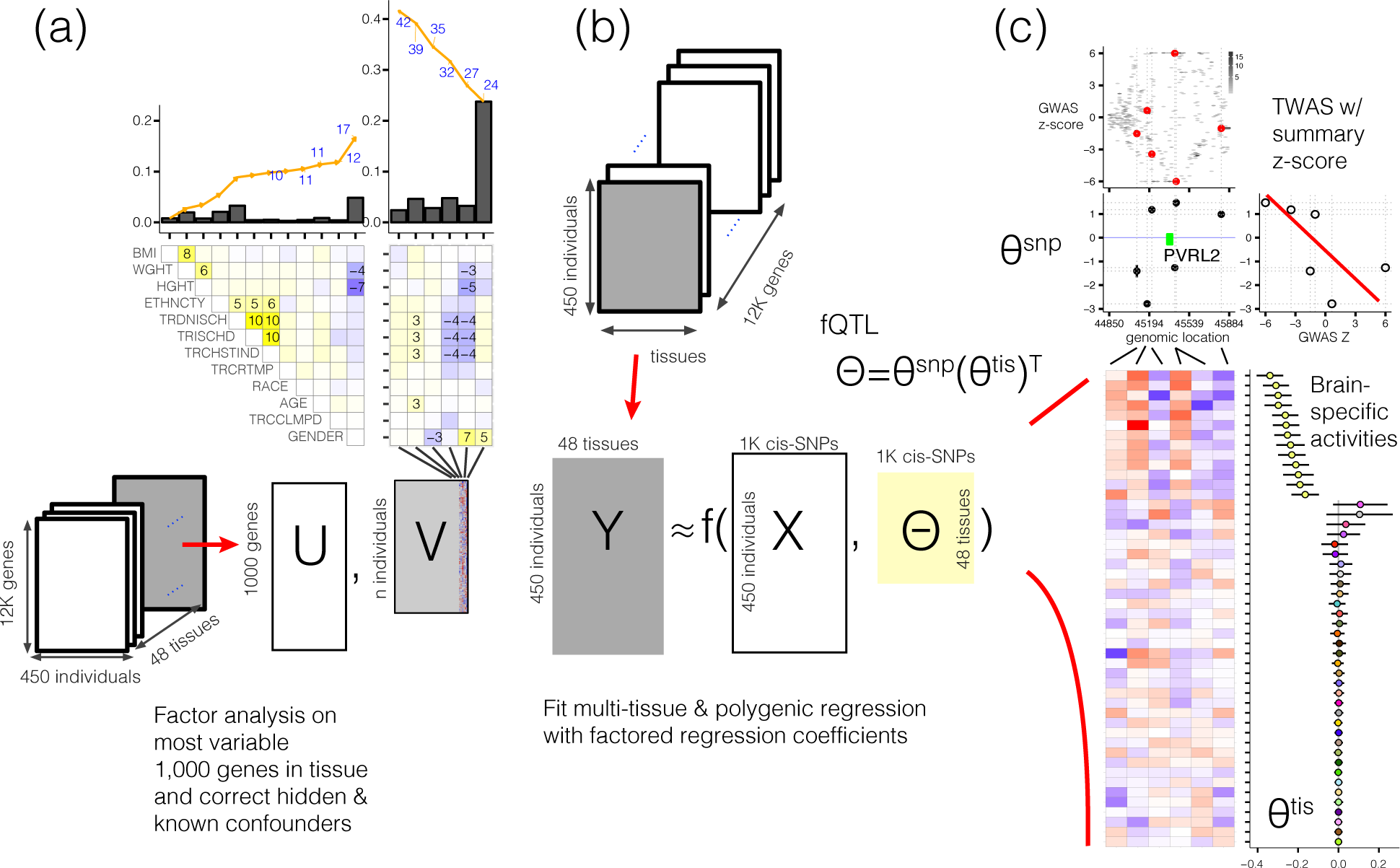
**Illustration of multi-tissue fQTL approach to identify tissue contexts in TWAS analysis.** (a) Sparse matrix factorization to learn and correct hidden confounders in each tissue. (b) Multi-tissue factored QTL (fQTL) estimation decomposing SNP-specific and tissue-specific effects. (c) TWAS with estimated SNP-specific effects and estimated tissue-specific context.

### B. Simulation study

We evaluated `fQTL` in realistic simulations and found it outperformed existing methods for a variety of scenarios. We estimated the statistical power for two inference problems: (1) identifying the causal SNPs and (2) identifying the tissues of action. We compared `fQTL` to the following methods:

- Single-tissue polygenic QTL models, which we refer to as `sQTL` (equivalent to fQTL with one tissue and *θ*^tis^ = 1)
- Lasso regression^29^, implemented in `glmnet` R package^30^ (`LASSO`). We tuned the regularization hyper-parameter λ by cross validation.
- Elastic-net^31^ regression, implemented in glmnet R package^30^ (ElasticNet). We tuned the regulariza - tion hyperparameter λ by cross validation, but we fixed the ridge-lasso tradeoff parameter to α = 0.5, which was used in previous TWAS analysis2.

We additionally sought to compare `fQTL` to the Bayesian sparse linear mixed effect model (BSLMM) implemented in `GEMMA`^17,32^. However, in preliminary experiements BSLMM could require an infeasible number of Markov Chain Monte Carlo iterations depending on the number of *cis*-SNPs. Furthermore, the convergence rate of the algorithm varied from gene to gene, so we did not include BSLMM in our analysis. In our experiments, when the model converged it performed equivalently to `Elastic-net` and `sQTL`.

A key problem in performing the proposed inference tasks is setting thresholds for discovery on the estimated model effect sizes. For causal variant prediction, we ranked SNP-tissue pairs according to the magnitude of regression coefficients |Θ_*jt*_| in decreasing order and calculated false discovery rate (FDR) and power using the PRROC package in R^33,34^. For tissue of action prediction, we calculated the proportion of variance explained (PVE) of gene expression by tissue-specific effects 
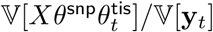
. We then ranked tissues in the decreasing order of PVE and calculated FDR and power.

We simulated multi-tissue expression datasets matching the level of missingness in GTEx using cis-genotypes in windows around randomly chosen genes varying the number of tissues of action, the number of causal eQTLs, and the heritability of gene expression. We first estimated the power to identify the causal SNP and found that fQTL had higher power except when there was only one tissue of action (Fig. 2a). In this case, fQTL pools information across irrelevant tissues, leading to poorer predictive models. Interestingly, as we increased the number of causal eQTLs (but still fixed a single tissue of action), fQTL power increased in causal variant identification, eventually matching the power achieved by Elastic-Net. As expected, as the number of tissues of action increases, fQTL outperforms single-tissue methods which are limited by sample size (missingness) in the training data. fQTL pools information across multiple tissues while favoring sparse solutions, allowing it to robustly identify causal SNPs without over-fitting and without pre-screening irrelevant tissues.

**Figure 2:**
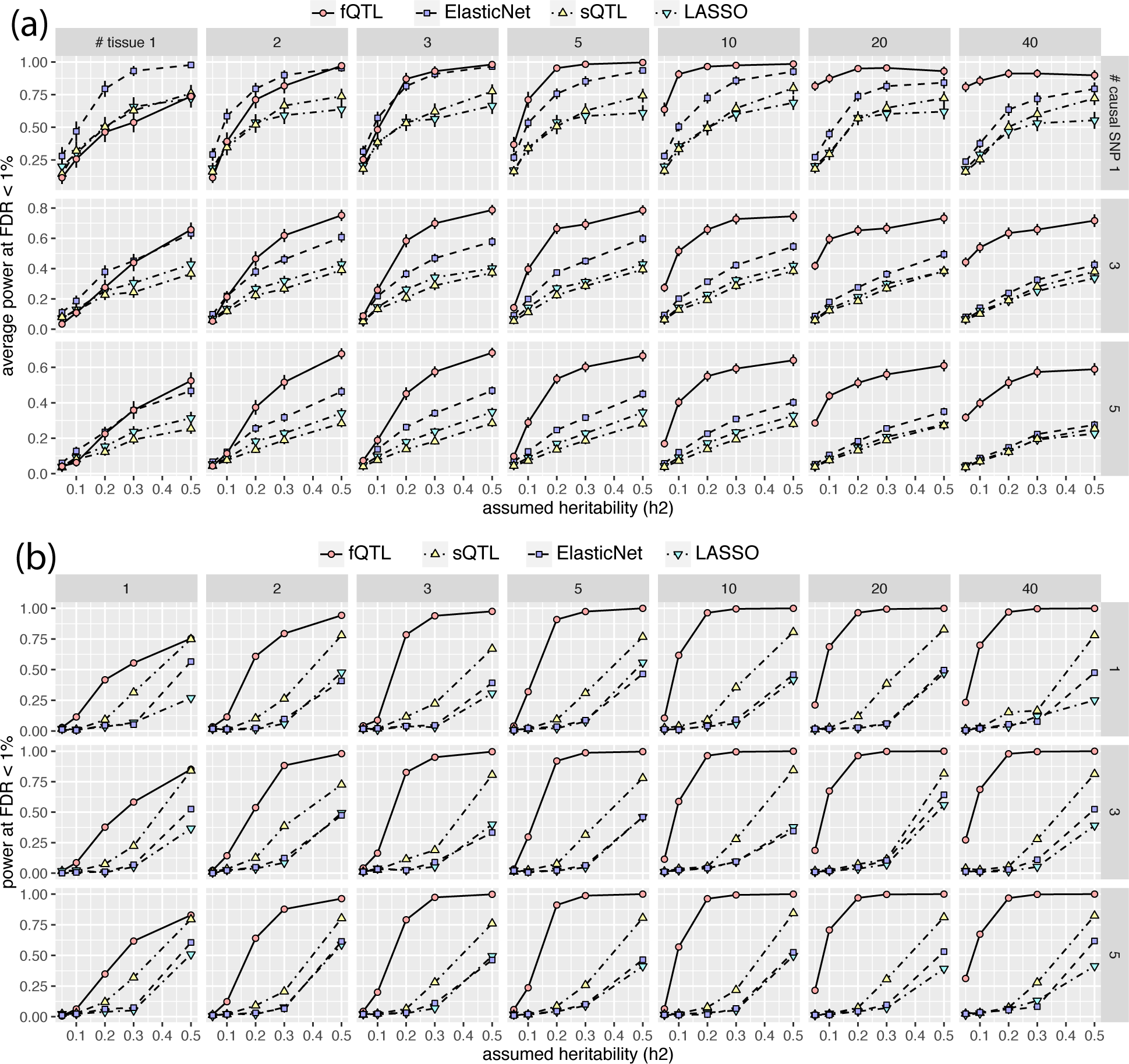
**Power calculation of different methods on simulated multi-tissue data.** Subpanels are arranged by different number of causal SNPs (rows; 1, 3, 5) and tissues (columns; 1 to 40). X-axis: assumed heritability. Y-axis: statistical power at FDR < 1%. (a) Power calculation of causal SNP prediction averaged over 175 repetitions. Error bars indicate 95% confidence intervals of the mean values (2 standard error). (b) Power calculation of tissue of action prediction.

We then investigated the power to detect the tissue of action (Fig. 2b). The results generally agree with those for causal variant identification. However, the number of causal SNPs had less impact on tissue prediction. Multi-tissue fQTL clearly dominated in all settings and gained more power as the number of tissues of action increased.

We finally investigated the sensitivity of our method to mis-specification of *K*. We first simulated scenarios where causal eQTLs were shared between *K* = 2 and *K* = 3 sets of tissues of action, but with different effect sizes in each subset (Supp. Fig. 7, *K* = 2; Supp. Fig. 8, *K* = 3). We found that fQTL assuming *K* = 1 performed only slightly worse than fQTL with the correct rank specification. In this study we did not explore learning *K* from the data because the GTEx data is too sparse (i.e., many tissues do not have enough individuals) to perform cross validation or hold-out validation. Importantly, fQTL with rank one still outperformed other single-tissue methods in these scenarios.

We then simulated a worst-case scenario for our model with arbitrary *K* by randomly assigning elements of Θ to be non-zero (Supp. Fig. 9). In this scenario, expression in different subsets of tissues of action is driven by different causal eQTLs. As expected, fQTL performed worse than single-tissue methods in identifying the causal variant, as it tries to pool information across irrelevant tissues. However, fQTL was still the most powerful method to identify the tissue of action when the SNP effect-mediated expression heritability *h*^2^ ≤ .3 and there were only few tissues of action. In this case, fQTL was able to capture some of the causal tissues and therefore successfully identify the causal eQTLs by pooling across those tissues.

### C. Identification of tissues of action in GTEx

Unlike our simulated data, significant PVE of real gene expression, such as that measured by the GTEx project, is attributable to experimental and technical confounding variables. For each gene in each tissue, we performed sparse matrix factorization to identify non-genetic, technical confounders on the 1000 most variable genes. We restricted our analysis to 48 tissues with sample size *n* ≥ 50, and found 2-15 non-genetic confounding factors in 40 of the 48 tissues. In the remaining 8 tissues, we found factors which were essentially 0. The learned factors explain a wide range of gene expression across tissues: for example, 15 factors explain 67% of whole blood gene expression variance, where 2 factors explain only 11% of spleen expression variance. Across the 48 tissues, 30% of variability is explained by non-genetic factors on average.

We found many of the inferred factors are highly correlated with known covariates: tissue ischemic time, age, body mass index, and gender (Supp. Fig. 10). However, in our analysis we included both known and learned covariates, again using the spike and slab prior to perform variable selection among the correlated covariates.

We next defined a set of 13,641 active genes (FDR 1%) by estimating the PVE of each gene’s expression explained by its *cis*-SNPs 
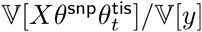
. We took the union of heritable genes identified across tissues. control the FDR of activ cted the null distribution of PVE by permuting the samples (Supp. Fig. 11). For comparison, we estimated the PVE explained by sQTL by fixing *θ*^tis^ = 1 and used the same procedure to identify active genes. As expected, the number of active genes per tissue was highly correlated with the sample size in that tissue (Fig. 3). However, fQTL increased power to detect active genes over sQTL, finding on average 3.8 times as many active genes across the 48 tissues.

**Figure 3:**
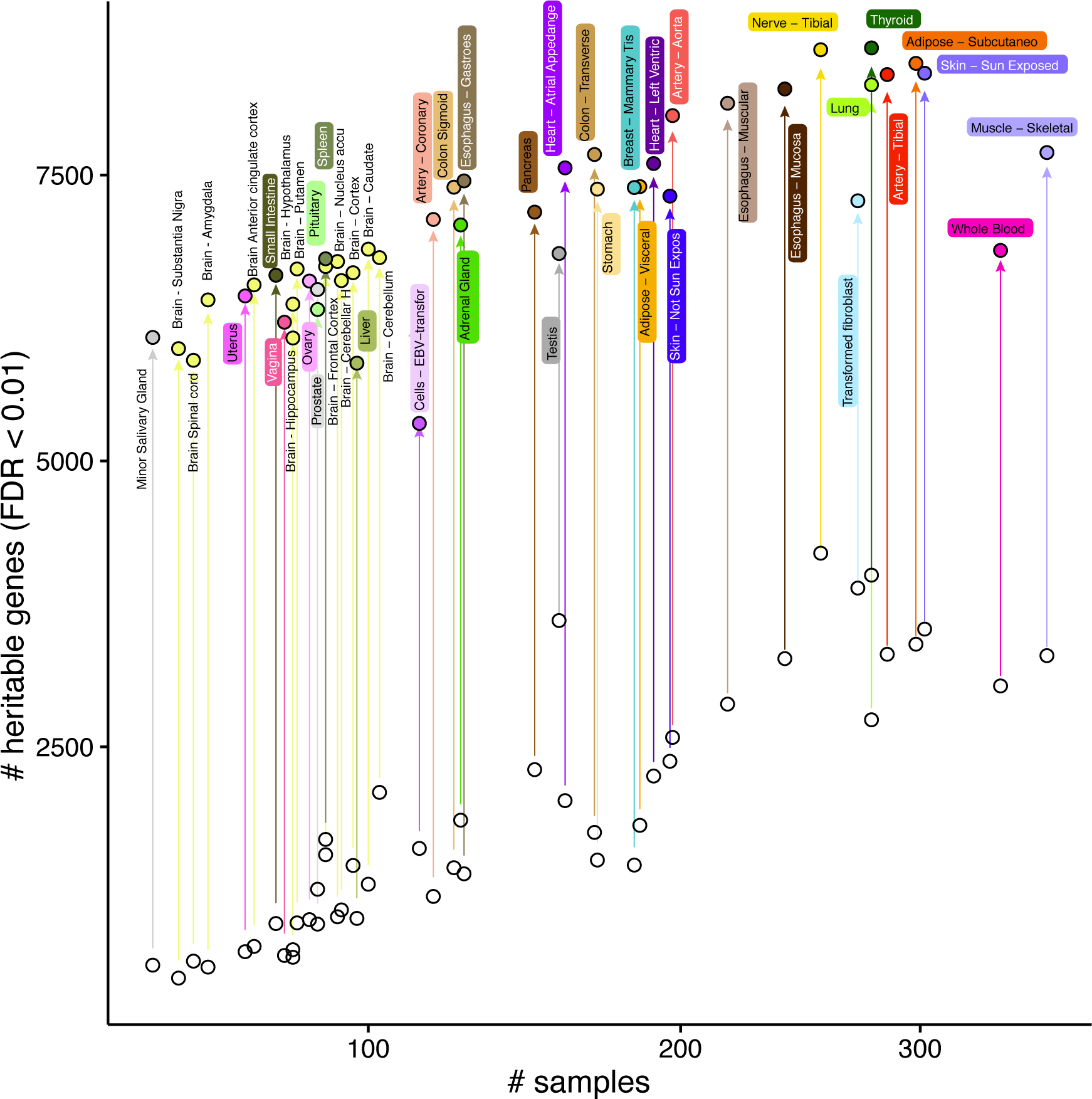
**Number of heritable genes across tissues.** Each tissue is represented as two points: empty circles denote the number of heritable genes within each tissue found by the single tissue polygenic model, where filled circles denote the number of heritable genes found by the multi-tissue polygenic model. Arrows match the points corresponding to the same tissue.

We then used the estimated tissue posterior inclusion probability 
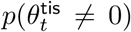
 to compute a correlation matrix between tissue activity patterns across all genes (Fig. 4). Our results broadly agree with those previously reported^35^: the brain tissues form a separate cluster, but non-brain tissues cluster together with no clear sub-clustering based on global tissue activity patterns. It is possible that non-brain tissues will form sub-clusters when examining tissue activity for subsets of genes in specific gene pathways; however, we do not explore that possibility in this study.

**Figure 4:**
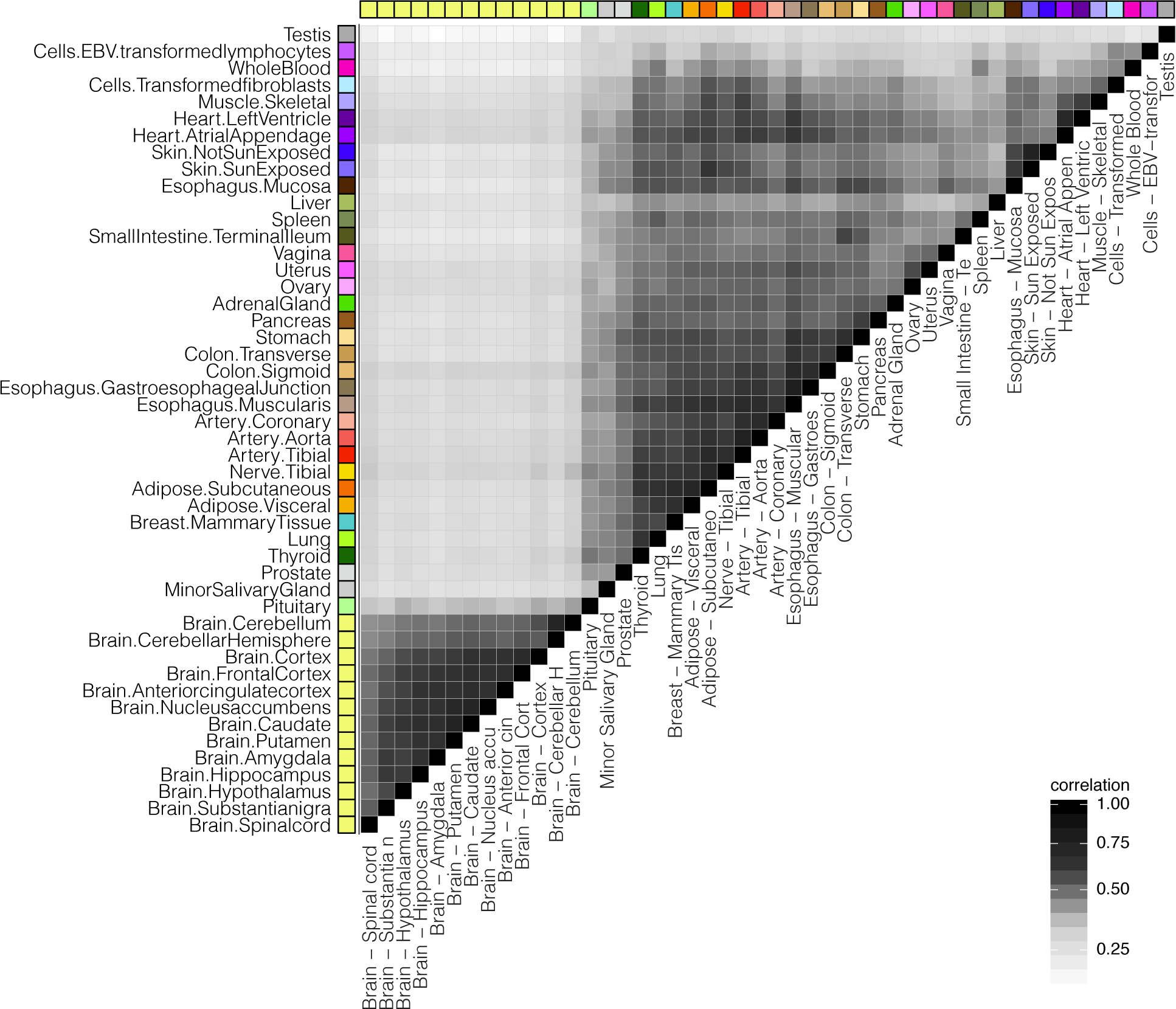
**Correlation between tissue activities in fQTL models.**

### D. TWAS using factored QTL effects

For each gene, we combined the estimated SNP and tissue effects from the fQTL model to estimate tissue-specific QTL effects and perform tissue-specific TWAS for two psychiatric disorders: Alzheimer’s disease^36^ (AD), and schizophrenia (SCZ)^37^. We have made all estimated fQTL model parameters and full TWAS summary statistics publicly available (https://github.com/ypark/fqtl).

We found 107 AD-associated genes at FDR 1% (lfdr calculation with normal null distribution by ashr package^38^; Fig. 5), of which only 10 have been previously reported in the NHGRI GWAS catalog^39^ as associated with AD. We found 40 genes significantly associated in brain tissues, of which 6 have been included in the GWAS catalog: BCAM, CR1, HLA-DRB1, HLA-DRB5, MRPL10, PVRL2.

**Figure 5:**
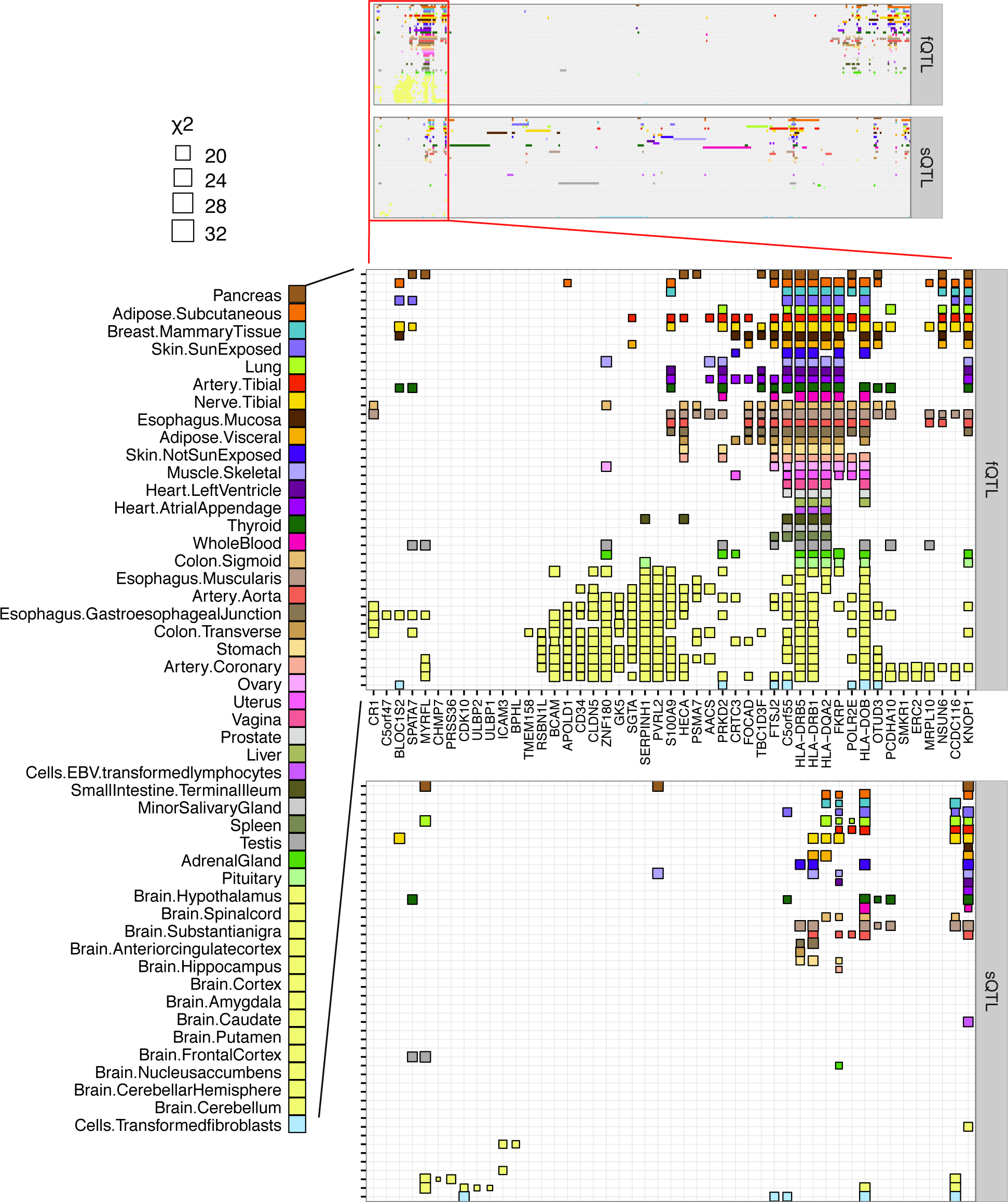
**Tissue-specific AD association statistics.** Columns include 346 genes significantly associated with AD at FDR 1% (107 fQTL; 282 sQTL); rows include 48 GTEx tissues. Sizes of blocks are proportional to χ^2^ statistics. Different colors indicate different tissue types.

We compared our tissue-specific TWAS results to single-tissue TWAS based on our sQTL models and found that fQTL greatly increased the power to detect both constitutively active and brain-specific genes associated with AD. Using sQTL, we found 282 AD-associated genes, including 15 known AD genes. However, 191 of the 282 genes are significant only in a single tissue, and only 11 are associated in brain tissues. In comparison, 5 of the 191 genes were found to be significant in more than one tissue by fQTL. None of the 15 single tissue TWAS genes in the GWAS catalog were significantly associated in brain, likely due to limited sample sizes. In comparison, 3 of these 15 genes were found by fQTL in brain tissues (PVRL2, HLA-DRB1, HLA-DRB5).

Two of the brain-specific AD TWAS genes are already known to be associated with AD. BCAM (basal cell adhesion molecule) and PVRL2 (poliovirus receptor-related 2) are both located in the APOE locus^40 –42^. However, BCAM is also confirmed as an AD-associated gene in follow up meta-analysis^43^. Similarly, PVRL2 remains significantly associated with AD after adjusting for LD structure^41^.

Our brain-specific TWAS identified SERPINH1 (Serine/Cysteine Proteinase Inhibitor 1), which has been previously associated with height in the GWAS catalog. SERPINH1 is elevated in microglial cells after treatment with amyloid beta, the classical hallmark of AD^44^. Differential expression of SERPINH1 is also linked with other psychiatric disorders such as major mood disorders^45^.

Our brain-specific TWAS also identified Claudin-5 (CLDN5), which is important for the function of the blood-brain barrier (BBB). In mice, brain injury followed by down-regulation of CLDN5 increased the permeability of BBB^46^. Depletion of CLDN5 in the blood-cerebrospinal fluid barrier may also lead to increased permeablility, imbalance of the immune system, and neurodegeneration^47^.

We found 382 tissue-specific TWAS genes in SCZ using fQTL (FDR 1%, Fig. 6), of which 45 have been previously reported in the GWAS catalog. 137 of the 382 genes are associated in brain tissues, of which 16 are in the GWAS catalog. As was the case for AD, we found that tissue-specific TWAS using fQTL greatly increased power to detect brain-specific TWAS genes. We found 845 single-tissue TWAS genes using sQTL, of which 82 have been previously reported in the GWAS catalog. However, 507 of the 845 genes are associated in only one tissue, only 61 genes of the 845 genes are significantly associated in brain tissues, and only 1 of the 61 brain TWAS genes (CYP2D6 associated in cerebellar hemisphere) is also included in the GWAS catalog.

**Figure 6:**
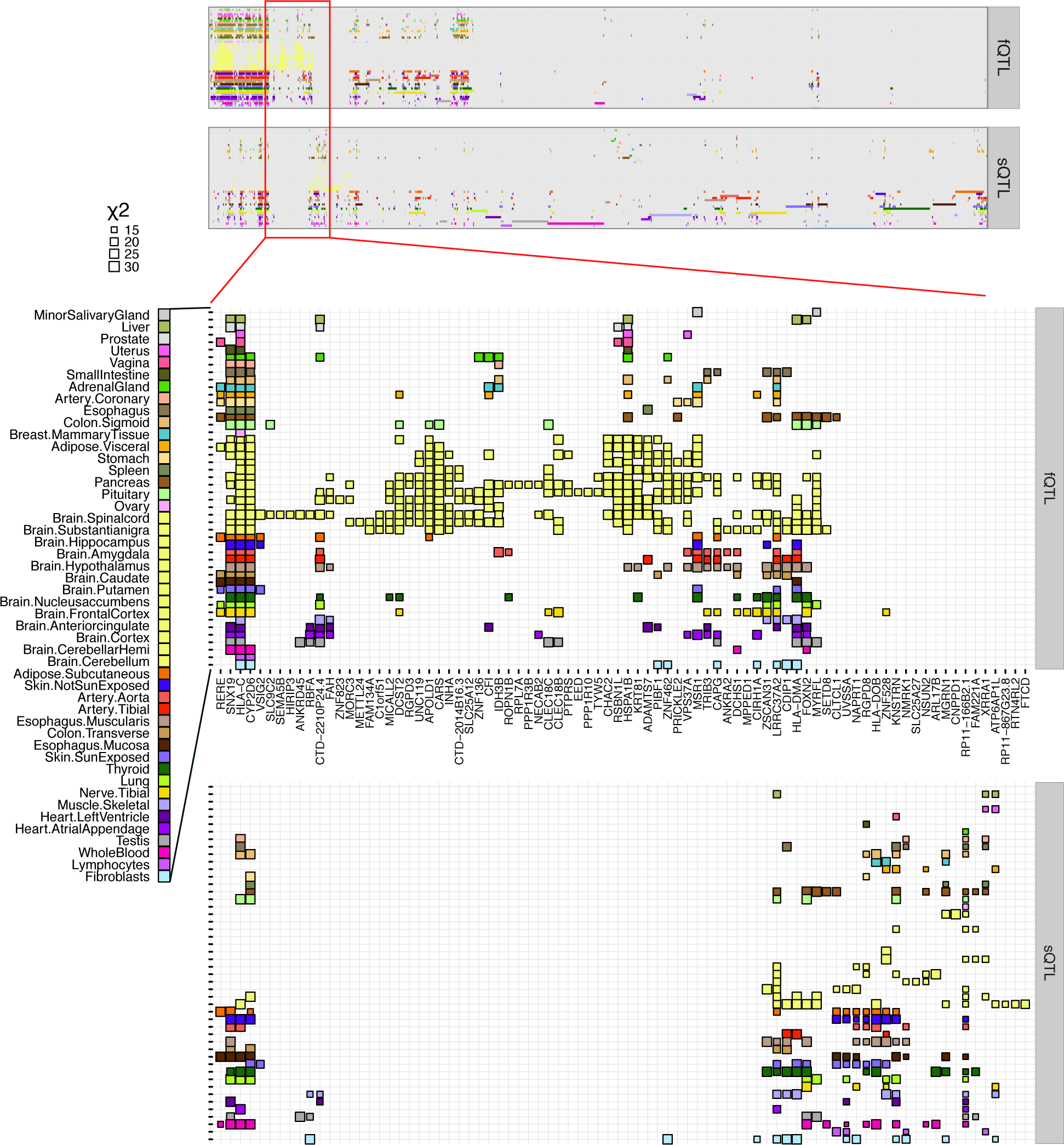
**Tissue-specific SCZ association statistics.** Columns include 1,007 genes significantly associated with AD at FDR 1% (368 fQTL; 845 sQTL); rows include 48 GTEx tissues. Sizes of blocks are proportional to χ^2^ statistics. Different colors indicate different tissue types.

We found a number of brain-specific SCZ TWAS genes which play known roles in other SCZ-relevant phenotypes: APOLD1, C1orf51 (CIART), CARS, and UNC119 (Supp. Fig. 14). APOLD1 (apolipoprotein L subunit D1), also known as VERGE (vascular early response gene), regulates vascular formation in the brain^48^ and is correlated with stress responses^49^.

C1ORF51 (circadian-associated transcriptional repressor; CIART) is important for circardian rhythm and is robustly expressed in the prefrontal cortex^50^. Circadian rhythms are disrupted in schizophrenia cases compared to controls^51^.

CARS (cysteinyl-tRNA synthetase) is associated with alcohol dependence and substance abuse^52,53^. Many reports document co-occurence of SCZ and substance abuse^54 –56^. UNC119 (uncoordinated 119) regulates an essential pathway of immunological synapse formation and T cell activation^57,58^. In mice, immature T cells trigger cognitive decline and behavioral abnormalities, but can be ameliorated by T-cell activation^59^.

## V. DISCUSSION

Here, we proposed fQTL, a multi-response multivariate method for eQTL mapping and tissue-specific expression prediction. We showed that our approach improves power over existing methods to detect both causal eQTLs and tissues of action by pooling information across tissues. We used our model to decompose tissue-invariant and tissue-specific QTL effects in the GTEx reference cohort, then combined these effects with summary statistics for AD and SCZ to perform tissue-specific TWAS and identify hundreds of new genes associated with these diseases.

There are a number of modeling and algorithmic improvements which can be made to our approach. Most importantly, in this study we assumed the eQTL-by-tissue effect matrix was rank one for intuitive interpretation: we fit a model with one vector of SNP-specific effects and one vector of tissue-specific effects. Our simulation revealed that mis-specification of this parameter leads to slight loss of power in our model. However, the problem of finding the optimal k requires some attention. We could determine *k* through cross-validation (although this is not possible in GTEx due to high missingness), optimizing the Bayesian information criterion^60^, or Bayesian model averaging over *K*.

We motivated our study of the rank one model by suggesting that the tissue-specific effect sizes of eQTLs observed in single-tissue approaches could be decomposed into tissue-invariant and tissue-specific components. In particular, we suggested that the tissue-specific component could be explained by variation across tissues in epigenomic state, or by altered expression of the upstream regulator in specific tissues. Distinguishing these two cases is necessary to interpret the mechanism of cis-regulation for genes identified by downstream TWAS and translate TWAS genes into therapeutics. However, the model we have proposed here cannot yet distinguish between these two cases. Future work should incorporate epigenomic information, transcription factor binding, and transcription factor expression to make specific testable biological predictions regarding transcriptional regulation of TWAS genes.

In this study, we assumed gene expression was Gaussian, and coerced the GTEx expression data to be approximately Gaussian. However, a more principled approach would be to use a more appropriate distribution (such as the negative binomial distribution) to directly model the process which generated the observed read counts. Existing methods cannot be easily extended to implement this approach because they make strong assumptions (like Gaussianity) and implement model-specific inference algorithms. Our SVI framework makes implementing extensions of this sort straightforward, without modification of the core inference algorithm. Future work should estimate the improvements in statistical power and false discovery rate when analyzing diverse data types in their observed, un-transformed distribution.

## ACKNOWLEDGMENTS

We thank Alexander Gusev, Benjamin Iriarte, Alkes Price, and Luke Ward for helpful discussions. We thank the International Genomics of Alzheimer’s Project (IGAP) for providing summary results data for these analyses. The investigators within IGAP contributed to the design and implementation of IGAP and/or provided data but did not participate in analysis or writing of this report. IGAP was made possible by the generous participation of the control subjects, the patients, and their families. The i-Select chips was funded by the French National Foundation on Alzheimer’s disease and related disorders. EADI was supported by the LABEX (laboratory of excellence program investment for the future) DISTALZ grant, Inserm, Institut Pasteur de Lille, Universite de Lille 2 and the Lille University Hospital. GERAD was supported by the Medical Research Council (Grant 503480), Alzheimer’s Research UK (Grant 503176), the Wellcome Trust (Grant 082604/2/07/Z) and German Federal Ministry of Education and Research (BMBF): Competence Network Dementia (CND) grant 01GI0102, 01GI0711, 01GI0420. CHARGE was partly supported by the NIH/NIA grant R01 AG033193 and the NIA AG081220 and AGES contract N01-AG-12100, the NHLBI grant R01 HL105756, the Icelandic Heart Association, and the Erasmus Medical Center and Erasmus University. ADGC was supported by the NIH/NIA grants: U01 AG032984, U24 AG021886, U01 AG016976, and the Alzheimer’s Association grant ADGC-10-196728.

## VIII. SUPPLEMENTARY METHODS

### A. Stochastic variational inference updates for sparse multivariate regression

Recall we assume the spike and slab prior on the parameter *θ*, and seek to find the optimal *q*(*θ*) to approximate the intractable posterior *p*(*θ* | *X*, *Y*,·). Under the variational approximation:

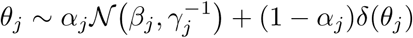

In order to find the optimal *q*, we need to optimize the evidence lower bound, which requires estimating a stochastic gradient using the log-derivative trick^21^:

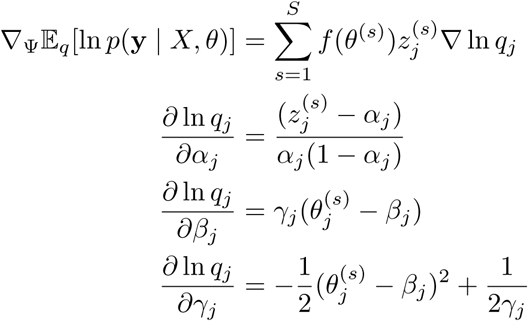

Using this approach, we can update (*α*, *β*, *γ*) by directly sampling *θ*^(*s*)^ ∼ *q*(*θ*), but this naive algorithm does not scale to high-dimensional models (*p* > 100 SNPs): sampling scales linearly in *p* and a large number of samples are required to accurately estimate the joint distribution of *θ*. In order to avoid sampling *θ*^(*s*)^ directly and speed up the inference, we re-parameterize:

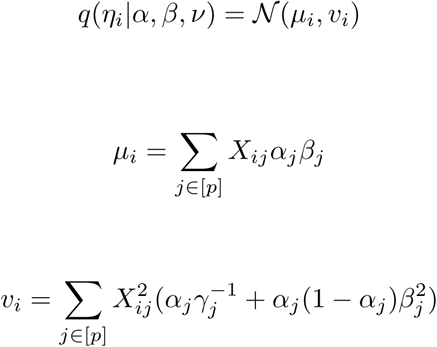

Now, we sample *ϵ*^(*s*)^ ∼ *N*(0, *I*), re-parameterize again to get a differentiable term^61^ 
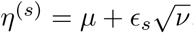
, and take gradients with respect to *μ*, *υ*:

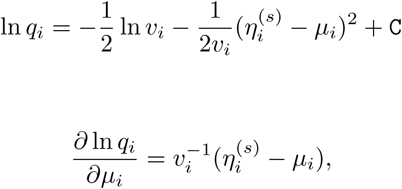

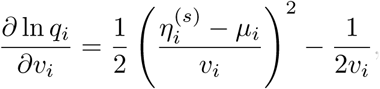

and with respect to *α*, *β*, *γ*: 
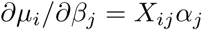
, 
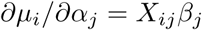
, 
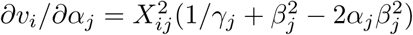
, 
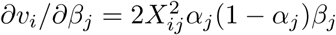
 and 
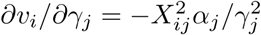
.

We can write the gradients using the chain rule, and efficiently compute them in parallel using matrix operations. For every stochastic vector *ϵ*^(*s*)^ we can evaluate log-likelihood vector f_*s*_, where 
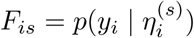
. First we can take derivatives with respect to 
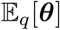
, in short 
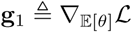
:

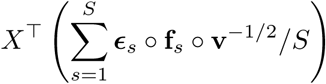

And with respect to 
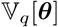
, in short 
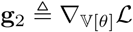
:

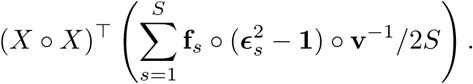

Applying the chain rule:

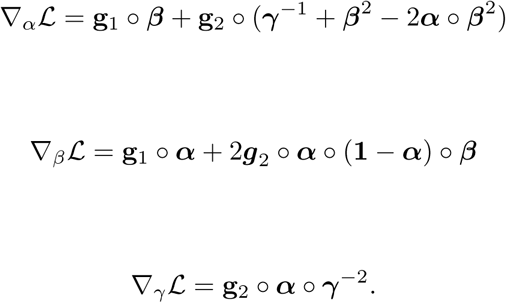

### B. Matrix factorization

For matrix factorization *H* = *UV*^T^ we used group spike-slab prior^18^:

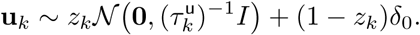

Under the variational approximation we define

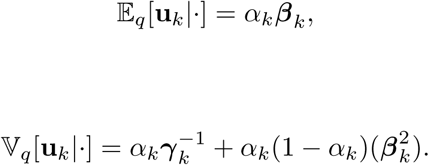

We use the same prior distribution and variational approximation for v_*k*_ vectors. We can characterize

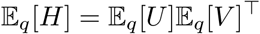

And

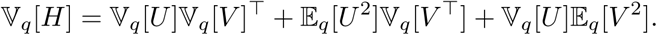

Sampling stochastic matrix *E*^(*s*)^,*E_ij_* ~ *N*(0, 1), we evaluate

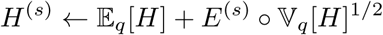

And log-likelihood *F*^(*s*)^ = ln *p*(*Y* |*H*^(*s*)^, ·) and estimate stochastic components of gradients:

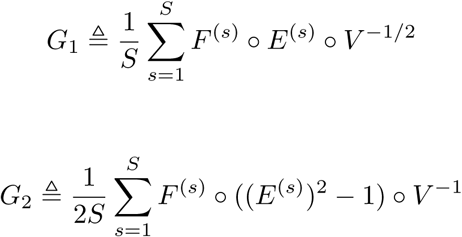

Using *G*_1_ and *G*_2_ we can derive gradient matrices straightforwardly

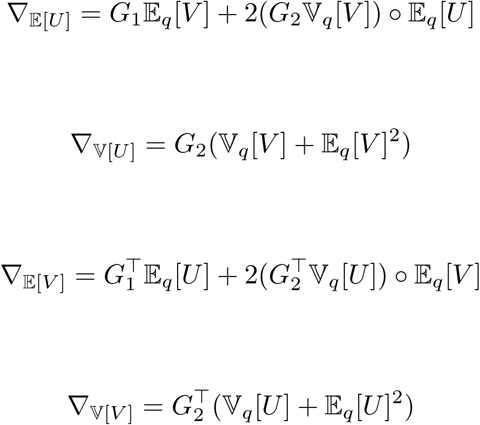

On these partial gradients we applying chain rules with respect to the original variational parameters.

### C. Control variate

Consider a generic variational approximation problem with surrogate distribution *q*(*θ* | λ) distribution *f*(*θ*). We want to calculate the gradient with respect to the variati for the intractable meter λ:

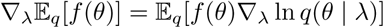

Define 
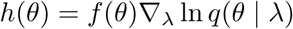
 and introduce a control variate *g*(*θ*) such that 
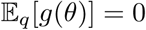
. Then, the gradient is equal to:

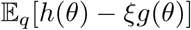

We can the find optimal ξ^*^ by solving:

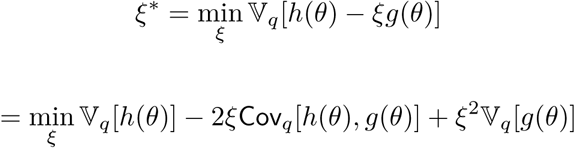

Previous work^21^ derived the optimal ξ:

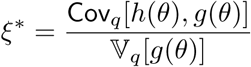

Here, we define 
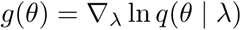
. Then, we have:

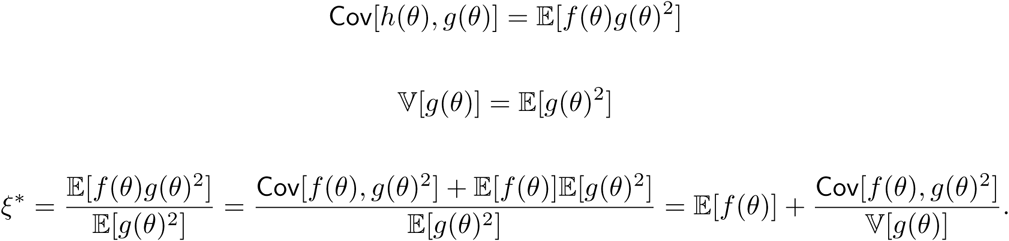

However, accurately estimating ξ^*^ requires many samples *θ*^(*s*)^ ~ *q*(*θ*^(*s*)^) | λ). Instead, we use 
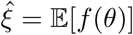
, resulting in a stochastic gradient estimator with variance:

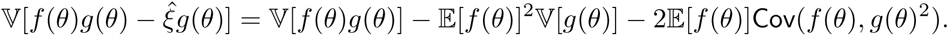

### D. Hyperparameter tuning

The spike and slab prior is defined by two hyperparameters, the prior inclusion probability *π* and prior effect size precision *τ*. In our extensive simulation experiments, direct optimization over the hyperparameters leads to degenerate solutions. To circumvent degeneracy, previous work used importance sampling or grid search^13,38^.

Here, we included the hyperparameters in the variational surrogate distribution. We re-parameterized:

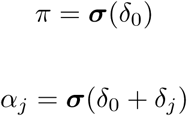

In our SVI updates, we updated *δ*_0_ and *δ*_*j*_ simultaneously. Similarly, for the prior precision *τ* and variational precision *γ_j_*:

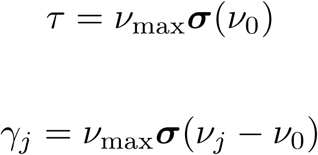

The model fit is highly sensitive to the setting of *ν*_max_. If the prior precision is too high, the prior dominates the likelihood and pulls effect sizes to zero. We set *ν*_max_ = 1000.

## IX. SUPPLEMENTARY FIGURES

**Figure 7:**
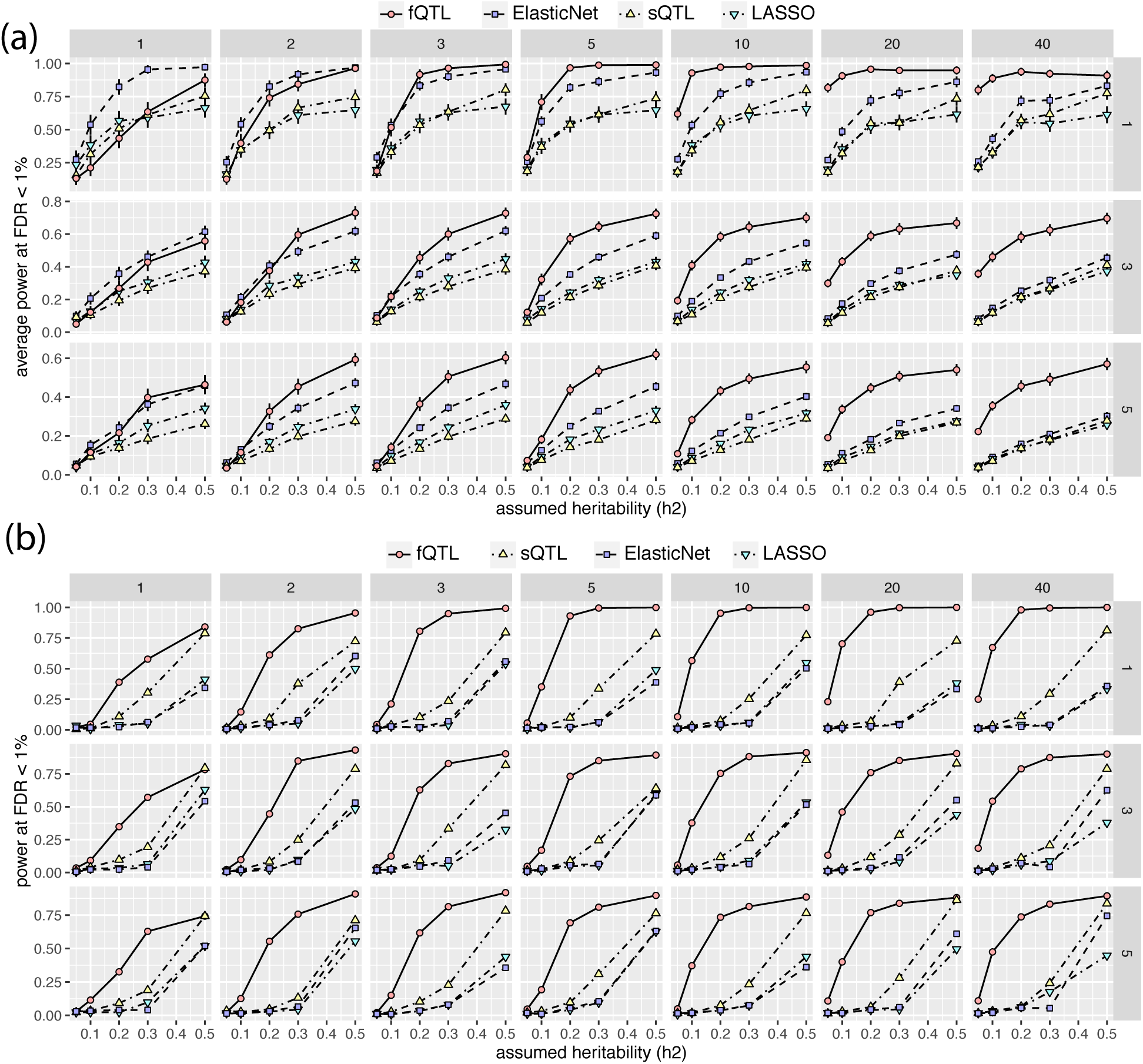
**Power calculation of different methods on multi-tissue data with mis-specified rank** *K* = 2. Subpanels are arranged by different number of causal SNPs (rows; 1, 3, 5) and tissues (columns; 1 to 40). X-axis: assumed heritability. Y-axis: statistical power at FDR < 1%. (a) Power calculation of causal SNP prediction. Error bars indicate 95% confidence intervals of the mean values (2 standard error). (b) Power calculation of tissue of action prediction.

**Figure 8:**
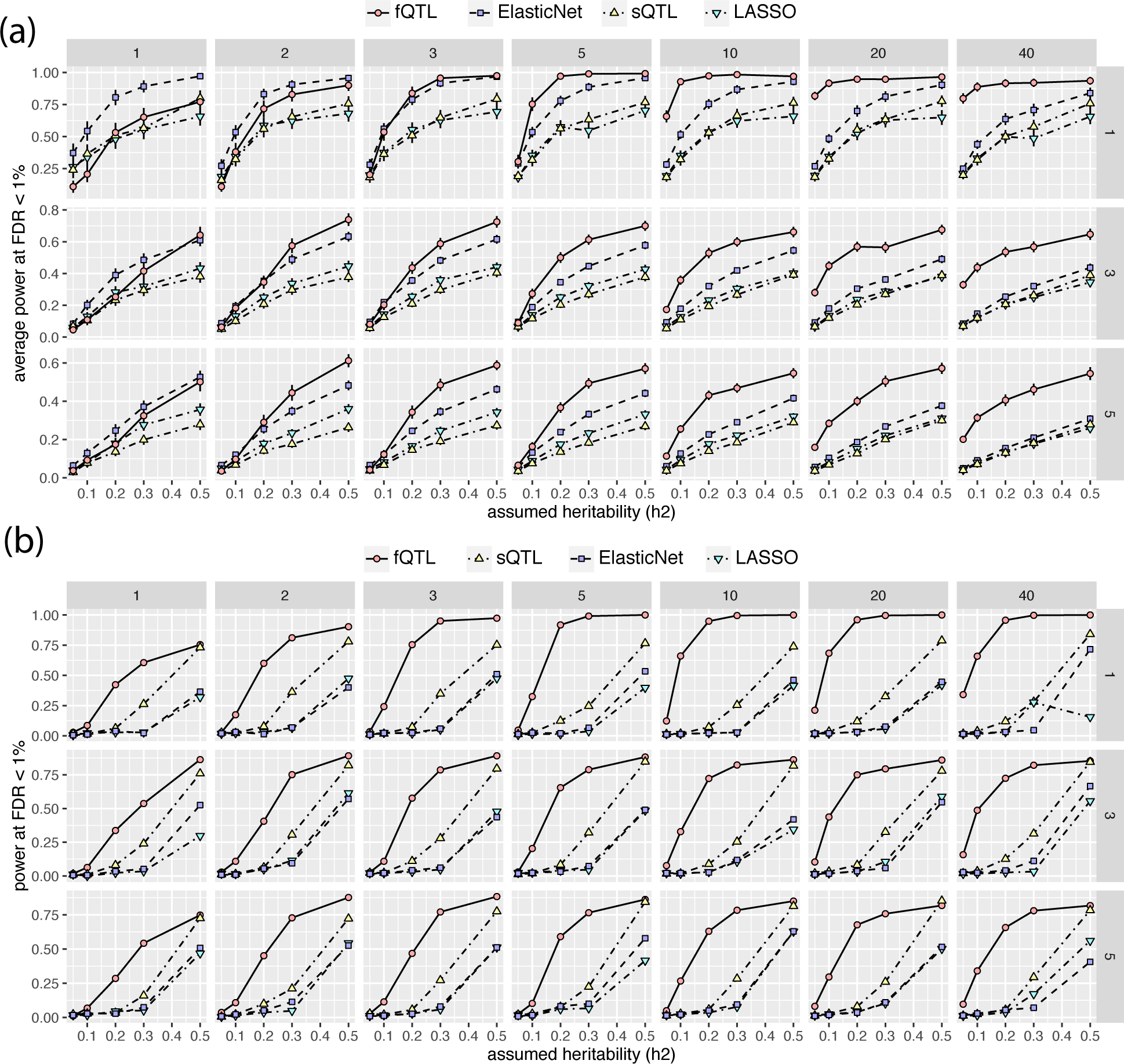
**Power calculation of different methods on multi-tissue data with mis-specified rank** *K* = 3. Subpanels are arranged by different number of causal SNPs (rows; 1, 3, 5) and tissues (columns; 1 to 40). X-axis: assumed heritability. Y-axis: statistical power at FDR < 1%. (a) Power calculation of causal SNP prediction. Error bars indicate 95% confidence intervals of the mean values (2 standard error). (b) Power calculation of tissue of action prediction.

**Figure 9:**
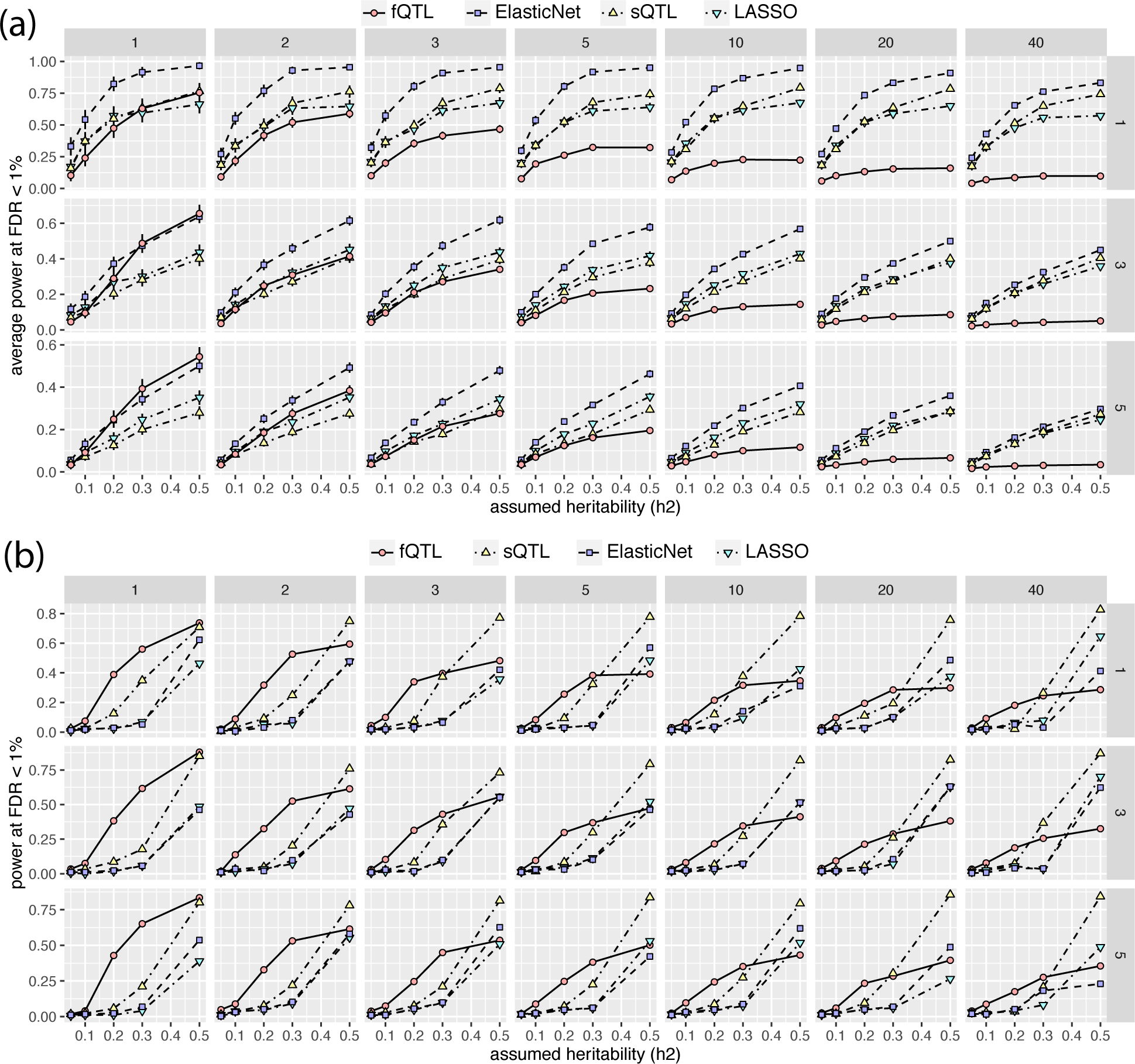
**Power calculation of different methods on unstructured multi-tissue data.** Subpanels are arranged by different number of causal SNPs (rows; 1, 3, 5) and tissues (columns; 1 to 40). X-axis: assumed heritability. Y-axis: statistical power at FDR & 1%. (a) Power calculation of causal SNP prediction. Error bars indicate 95% confidence intervals of the mean values (2 standard error). (b) Power calculation of tissue of action prediction.

**Figure 10:**
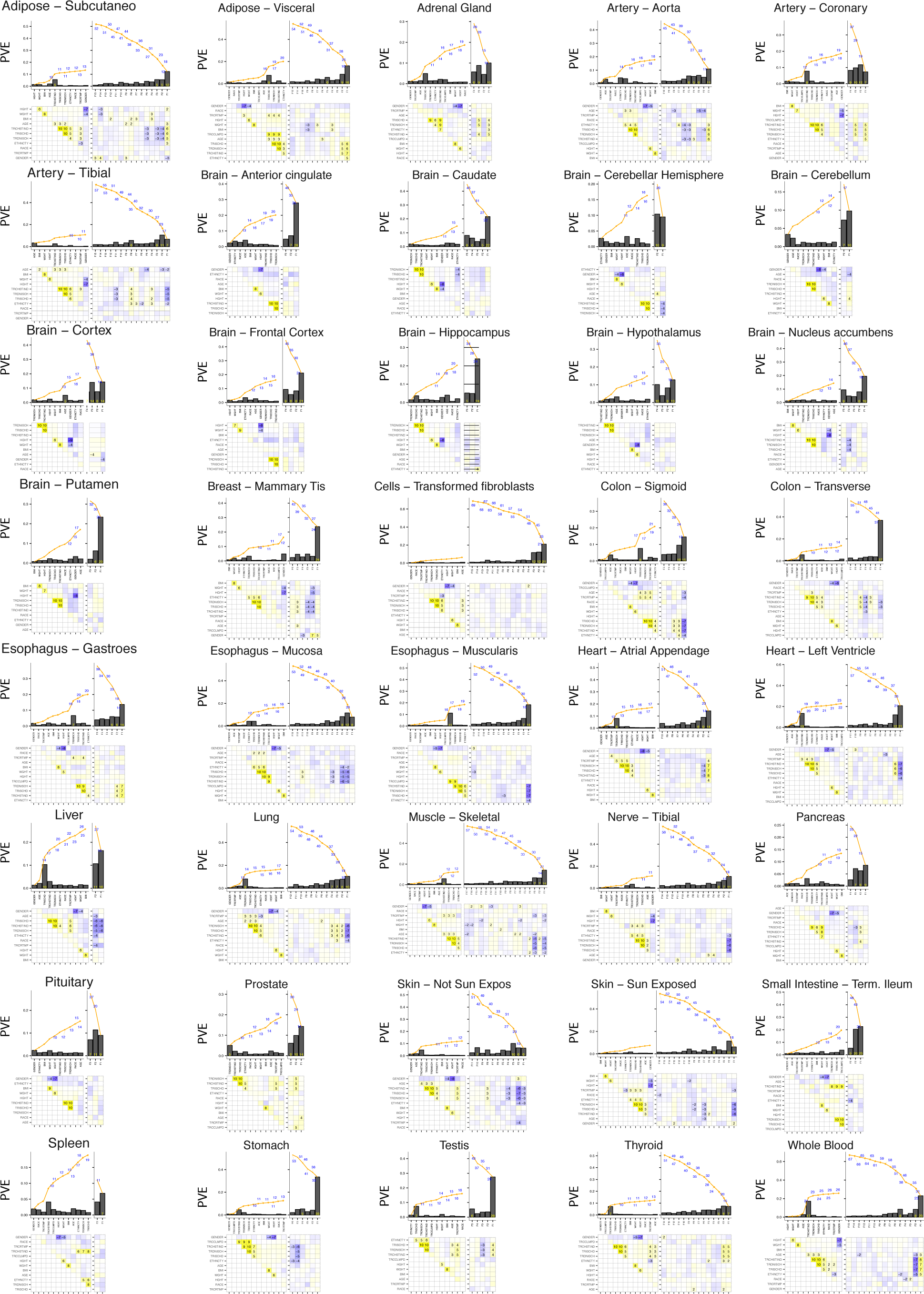
**Association of tissue-specific hidden factors with known variables.**

**Figure 11:**
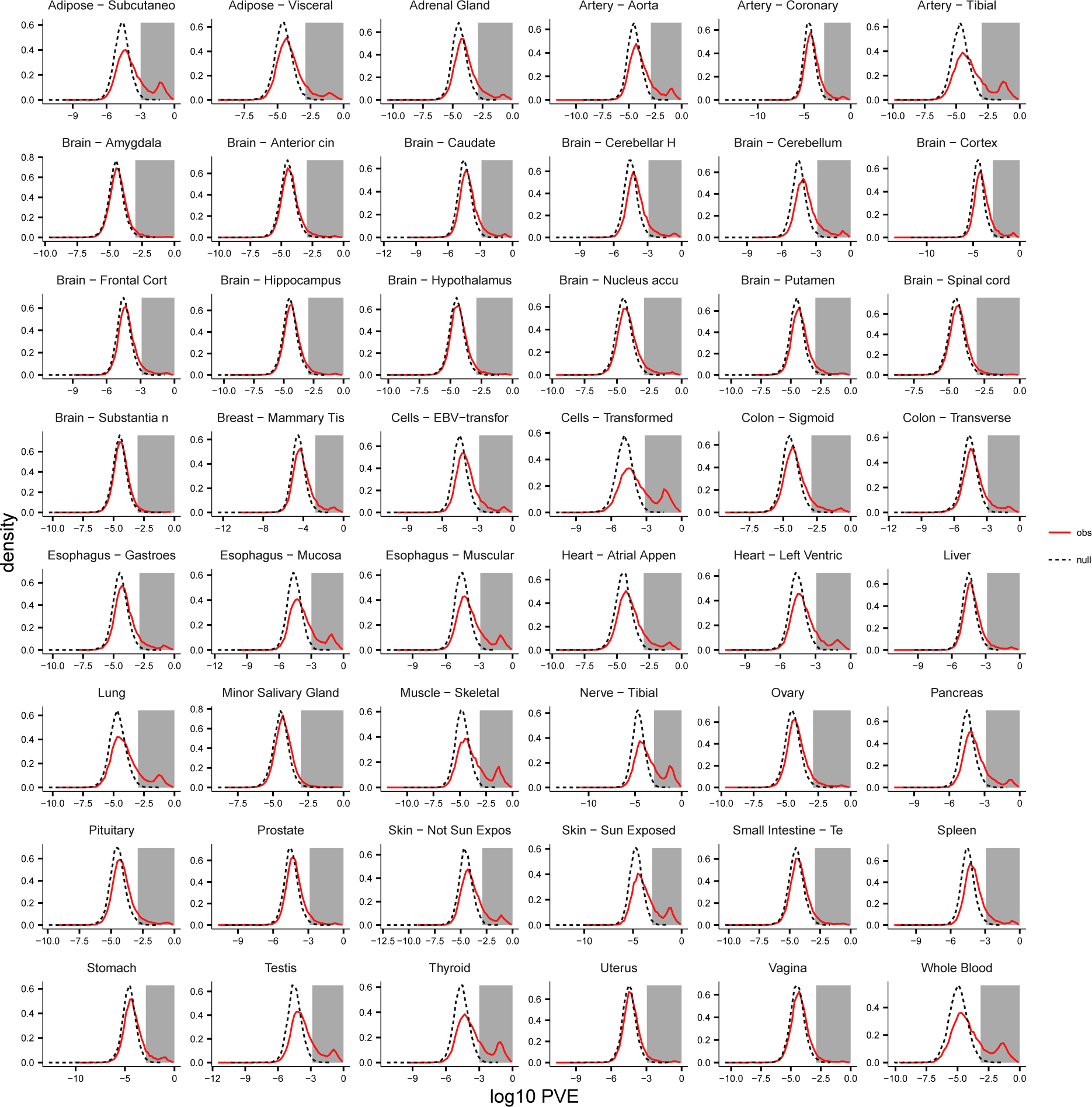
**Tissue-specific null distribution sQTL PVE constructed by sample permutation.**

**Figure 12:**
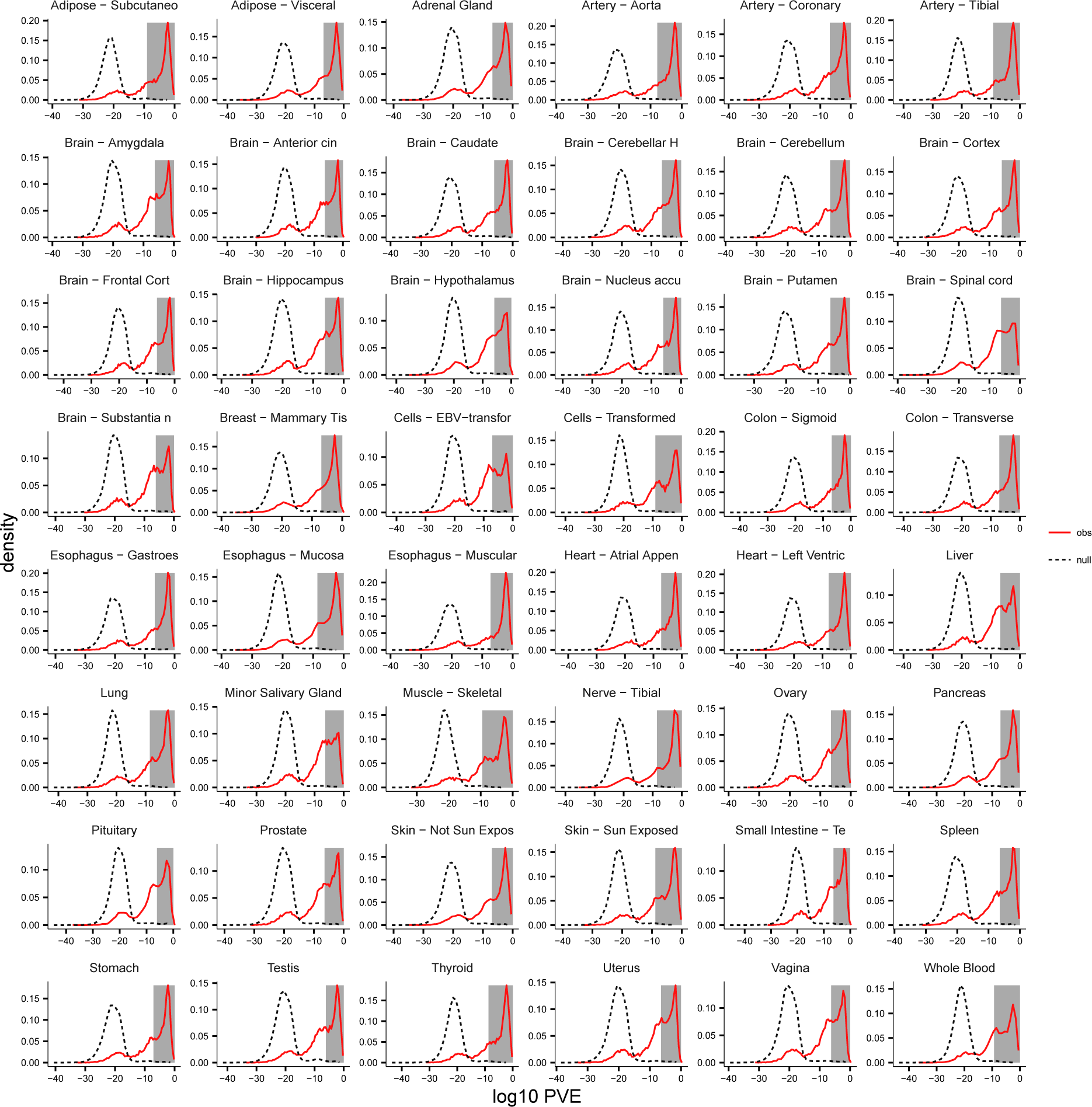
**Tissue-specific null distribution fQTL PVE constructed by sample permutation.**

**Figure 13:**
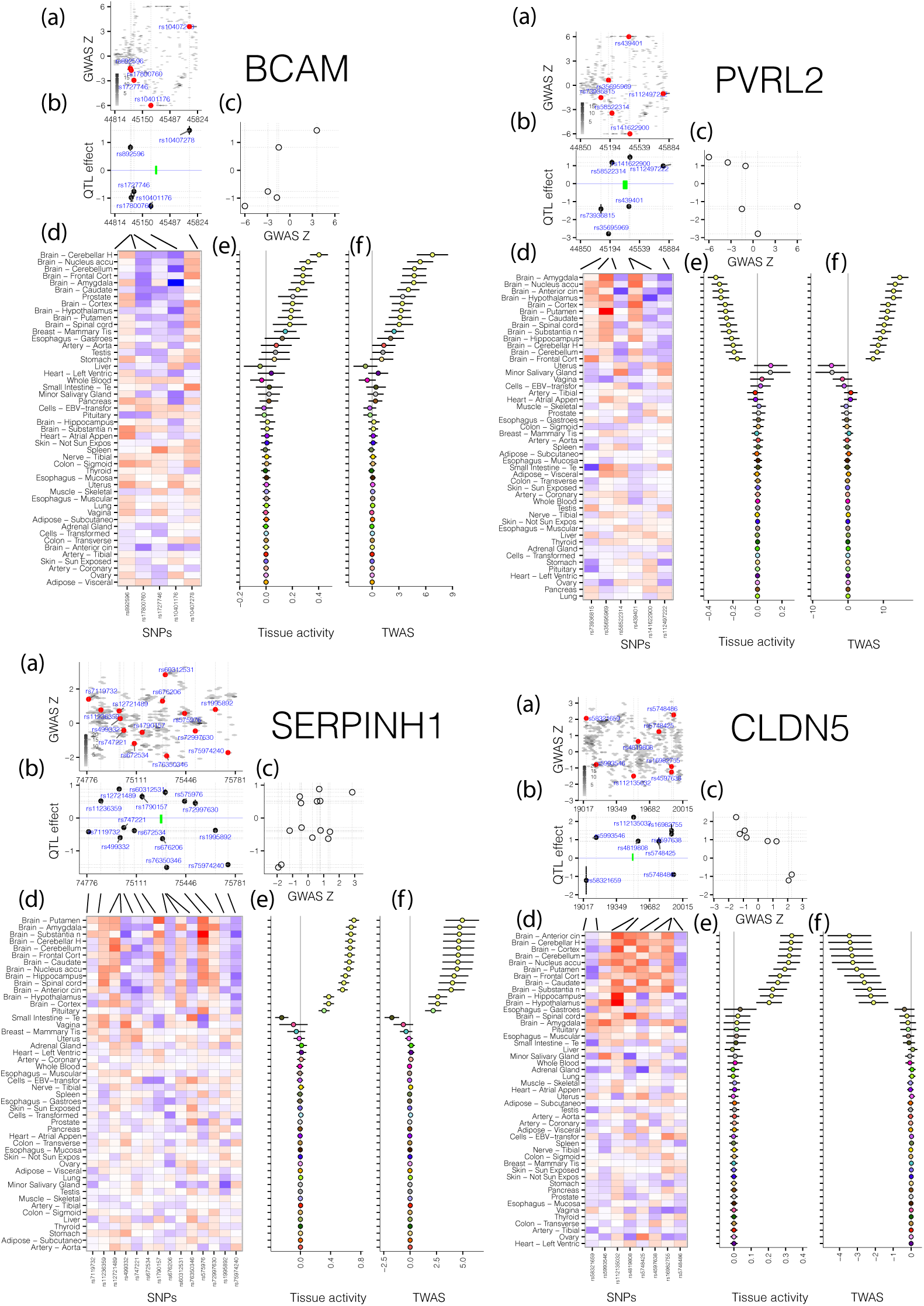
**Examples of brain-specific AD genes.** (a) Distribution of GWAS z-scores in the *cis*-region. (b) Polygenic QTL effect sizes in the same region. (c) Correlation between QTL and z-score vectors. (d) Marginal correlation matrix of SNP-tissue pairs. (e) Tissue activity effects found by fQTL model. (f) TWAS statistics. Errorbars denote 95% confidence intervals.

**Figure 14:**
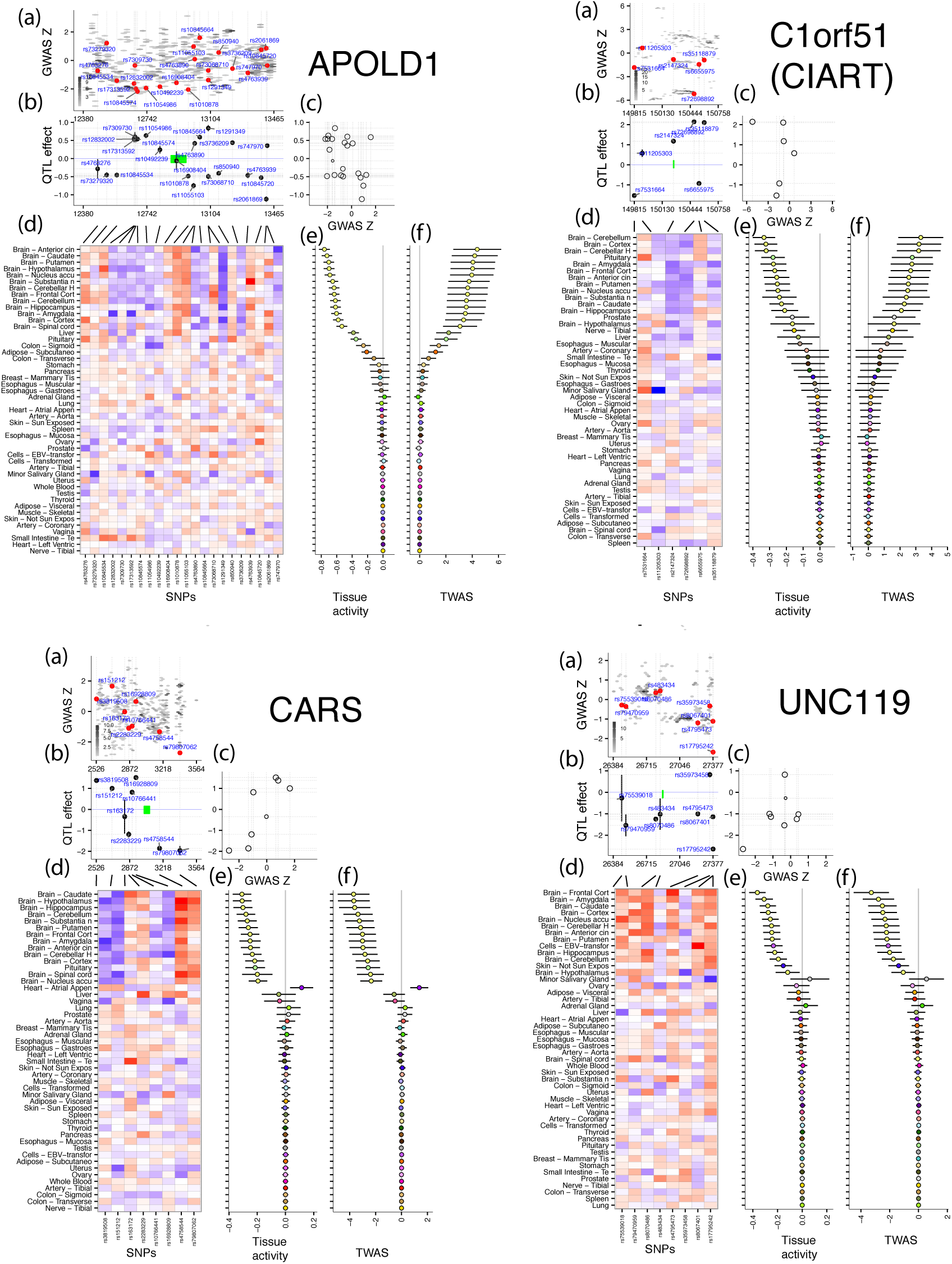
**Examples of brain-specific SCZ genes.** (a) Distribution of GWAS z-scores in the *cis*-region. (b) Polygenic QTL effect sizes in the same region. (c) Correlation between QTL and z-score vectors. (d) Marginal correlation matrix of SNP-tissue pairs. (e) Tissue activity effects found by fQTL model. (f) TWAS statistics. Errorbars denote 95% confidence intervals.

## X. SUPPLEMENTARY TABLES

Full gzipped text files on TWAS results are made available at https://github.com/YPARK/FQTL/tree/master/GTEx.

